# Direction-dependent bias in the perceived approach speed and its implications for robot motion design

**DOI:** 10.64898/2026.01.15.699393

**Authors:** Tatsuto Yamauchi, Hideki Tamura, Tetsuto Minami, Shigeki Nakauchi

## Abstract

Humans frequently interact with robots that approach from different directions; however, how the approach direction systematically biases the perceived speed remains unclear. Here, we demonstrate a robust direction-dependent bias in the perceived approach speed, whereby objects approaching from behind are consistently perceived as moving faster than those approaching from the front are. Across five experiments using both physical robots and immersive virtual reality, this bias manifests as a systematic shift in the point of subjective equality without corresponding changes in discrimination sensitivity, indicating a perceptual bias rather than altered precision. We further show that this effect can be explained by body-induced differences in interocular velocity that arise when observers orient themselves toward stimuli approaching from behind. This bias persists in virtual environments and for nonrobotic objects, suggesting that it is driven primarily by visual factors rather than multisensory cues or robot-specific appearance. Together, these findings provide a perceptual foundation for incorporating approach direction into the design and evaluation of the motion behavior of robots in human-centered environments.

## I. INTRODUCTION

In recent years, the deployment of autonomous mobile robots (AMRs) has rapidly expanded in urban environments, resulting in increasingly frequent human–robot interactions (HRIs) in domains such as logistics, hospitality, and guidance support. Beyond functional performance, achieving smooth social acceptance requires robot designs and control strategies that account for fundamental characteristics of human behavior. For example, Kamezaki proposed a control method in which a robot adaptively selects between assertive path planning and yielding on the basis of its relative position with respect to a human, demonstrating that humans prefer reciprocal interactions in which both agents adjust their trajectories rather than unilateral yielding by the robot [1]. Similarly, model-based obstacle avoidance algorithms that incorporate human motion, intentions, and preferences have been shown to improve interaction quality [2]. Consistent with these findings, humans generally tend to yield to AMRs during spatial interactions [3], and robots employing human-like avoidance trajectories or pedestrian models elicit greater psychological comfort than those relying on conventional algorithms do [4], [5]. Humans also prefer proactive assistance, in which an AMR approaches an object to support a task rather than passively waiting, as observed in luggage transport scenarios [6], although close proximity can reduce the walking speed and increase hesitation. Moreover, human responses to robots are modulated by social cues such as facial expressions and attitudes [7], highlighting the importance of optimizing a robot’s appearance and communication modalities to facilitate the acceptance of its intentions [8]. Collectively, these studies highlight the need to ground robot behavior design in a principled understanding of human behavioral traits in shared environments.

Building on this human-centered perspective on robot behavior design, a growing body of research has shown that a robot’s approach behavior plays a critical role in shaping human comfort, arousal, and perceived safety. In particular, approach speed and acceleration have been reported to modulate arousal and threat perception [9], with slower approaches generally perceived as being safer [10]. However, uniformly reducing robot speed is not a viable solution, as excessively low velocities can disrupt pedestrian flow and compromise efficiency in shared environments [11]. This limitation highlights the need for the adaptive modulation of approach behavior on the basis of contextual factors. In realistic settings, robots frequently encounter humans from various directions, and prior studies have demonstrated that acceptable interpersonal distance depends on the approach direction [12]. Moreover, perception in the rear space exhibits cognitive asymmetries relative to the frontal visual field [13], and being approached from behind by a robot has been shown to elicit greater discomfort than frontal encounters do [14]. Together, these findings suggest that approach direction is a key but underexplored factor that may critically modulate how robot motion is perceived.

Despite these advances, research on speed perception has focused almost exclusively on objects approaching the frontal visual field [15], [16], [17]. Consequently, whether—and in what manner—perceived approach speed differs between front and behind-approaching objects remains unclear. This limitation is particularly important given accumulating evidence that human perception exhibits systematic asymmetries between the front and behind space across multiple sensory modalities, including vision and audition [13], [18], [19]. Although such front-behind asymmetries have been reported across modalities, the present study focuses on vision, as visual geometry is directly constrained by head and body orientation during rearward observation. If such perceptual asymmetries are rooted in the bodily and geometric constraints of the human observer, a plausible contributing factor is the asymmetric visual input that arises when orienting toward stimuli approaching from behind, where imperfect alignment of the head and torso leads to asymmetric visual geometry across the two eyes and may give rise to interocular velocity differences (IOVDs). Although IOVDs provide a potential mechanistic account linking body-induced asymmetries to visual motion processing, their role in direction-dependent speed perception has not been systematically examined, particularly in the context of human– robot interactions.

In this study, we investigate whether perceived approach speed differs between objects approaching from the front and those approaching from behind and whether any such directional bias can be attributed to body-induced visual asymmetry. We hypothesize that approaches from behind are perceived as being faster than frontal approaches are, independent of learning direction or stimulus appearance. To test this hypothesis, we conducted a series of five experiments. Experiments 1–3 examine the existence and nature of perceptual bias, as well as its underlying visual mechanism, using physical robots. In addition, experiments 4 and 5 further assess the robustness and generality of the effect in immersive virtual environments and with nonrobotic objects.

## II. Methods of Experiments 1 & 2

### A. Participants

All the participants were students at Toyohashi University of Technology. The target sample size was set at 30 participants per experiment to exceed the minimum requirement indicated by an a priori power analysis, which yielded a required sample size of 29, assuming a statistical power of 0.8, an alpha level of 0.05, and an effect size of Cohen’s *d* = 0.54 estimated from a pilot experiment (*n* = 4). Accordingly, Experiments 1 and 2 each involved 30 participants (Experiment 1: 7 females, 23 males; mean age = 22.3 ± 0.8 years; Experiment 2: 6 females, 24 males; mean age = 21.6 ± 1.7 years).

Across all five experiments, the experimental protocols involving human participants were approved by the institutional review board of Toyohashi University of Technology (approval number: 2024-28). Written informed consent was obtained from all the participants prior to participation.

### B. Apparatus

A custom wheeled platform (Mega Rover Ver. 3.0, Vstone) was used as the AMR (Fig. 1a), which measured 330 mm × 270 mm × 800 mm (length × width × height). Two motors connected to the wheel axles were controlled by an onboard computer. A separate computer located in the experimental chamber received position data from a tracking system (described below) and transmitted control signals to the onboard computer via Wi-Fi. Both computers were operated using the Robot Operating System 2 (ROS 2) Humble running on Ubuntu 22.04 LTS. The AMR was equipped with a motion tracker (VIVE Tracker 3.0, HTC) mounted on its front.

**Fig. 1.**
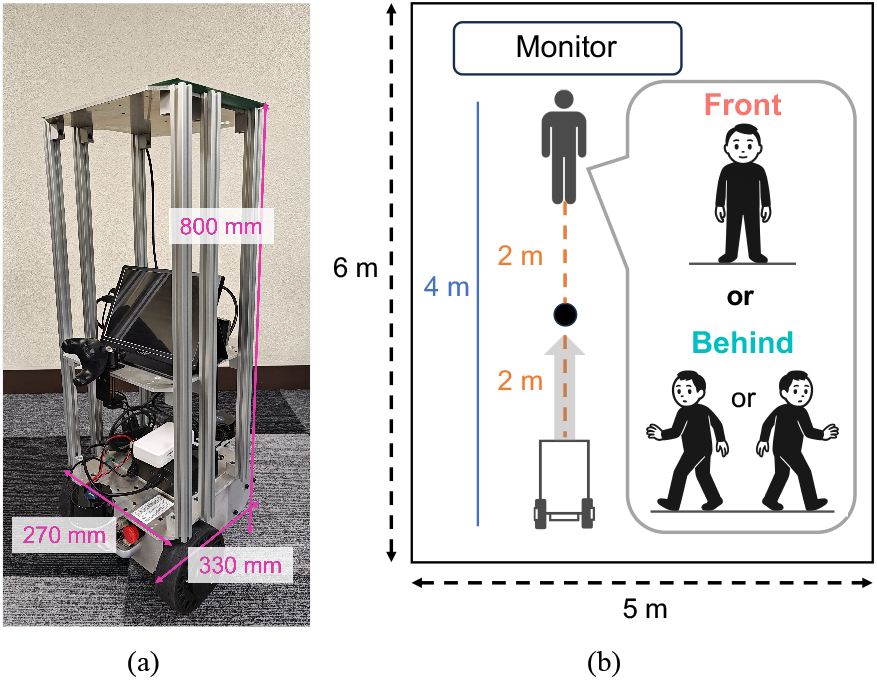
Experimental setup. (a) Autonomous mobile robot in the experiment. (b) Illustration of the environment.

The motion tracking system, running on Windows 11, communicated with the trackers via Unity (2022.3.21f1), SteamVR (version 2.5.5), and the SteamVR Unity Plugin (version 2.8.0; SDK 2.0.10). The system recorded the position and orientation of each tracked body part at a sampling rate of 90 Hz within a play area defined by four base stations (SteamVR Base Station 2.0) installed in the experimental room.

To monitor whether participants were facing forward or backward, a motion tracker was affixed to their waist. The participants also wore a head-mounted eye tracker (Pupil Labs Neon) to record their point of gaze. The experimental environment is illustrated in Fig. 1b. The experimental area measured 5 m × 6 m (width × length). The AMR initiated its approach 4 m from the participant and stopped at 2 m. Only one participant and one AMR were present in the experimental field at any given time, and the experimenter operated the control and tracking systems from outside the field.

### C. Procedure

Each experiment employed a bisection method consisting of two phases: a speed memorization phase and a main task. During the speed memorization phase, participants learned two reference approach speeds of the robot (slow: 0.5 m/s; fast: 1.0 m/s) while maintaining a fixed viewing direction—facing forward in Experiment 1 and facing backward in Experiment 2. In this phase, the AMR approached at either low or high reference speed in a random order, and the participants indicated whether the perceived speed was low or high. Trials were repeated until participants achieved five consecutive correct responses for each reference speed.

In the main task, participants judged whether the AMR’s approach speed was closer to the low or high reference speed. The AMR approached at one of seven speeds (0.5, 0.6, 0.7, 0.75, 0.8, 0.9, and 1.0 m/s) in a random order. A total of 112 trials were conducted (7 speeds × 2 approach directions [front/behind] × 8 repetitions), with the trial order being randomized separately for each participant. The trials were divided into four sessions, and the participants were allowed to take an optional break between sessions. After half of the sessions were completed, the participants repeated the speed memorization phase, with a reduced criterion of three correct responses for each reference speed.

Before the start of each trial, the AMR returned to its initial position, which was 4 m from the participant. During this repositioning period, participants faced a monitor positioned opposite the AMR and waited for the next instruction. This procedure ensured that participants did not observe the AMR while it was returning to the starting point, thereby preventing the return motion from influencing subsequent speed judgments. Once the AMR reached the initial position, the monitor displayed one of three posture images corresponding to the approach direction condition. These conditions included one front condition and two behind conditions, indicating turning left or right to look behind. In the front condition, participants oriented their body to face the AMR, whereas in the behind condition, participants rotated their upper body to look behind. The AMR then emitted an auditory beep to signal trial onset and approached the participant, stopping at a distance of 2 m. After the AMR stopped, participants responded using a keypad, pressing the “+” key (labeled “Fast” on the screen) if the approach speed was closer to the high reference speed or the “–” key (labeled “Slow”) if the approach speed was closer to the low reference speed.

Prior to the experiment, participants received detailed instructions regarding the task procedures, trial sequence, and total number of trials. Participants were additionally instructed to follow specific behavioral constraints: (1) rotate only their upper body while keeping the lower body stationary when looking behind, (2) fix their gaze on the motion tracker mounted at the center of the AMR, and (3) hold the response keypad in their hands throughout the experiment. No further instructions that could bias participants’ behavior were provided.

### D. Data analysis

Response data were analyzed by fitting the probability of “fast” responses at each speed level with a psychometric function using a generalized linear model with a probit link function implemented in R (version 4.5.0). For each participant, the point of subjective equality (PSE) and the just noticeable difference (JND) were estimated from the fitted psychometric function. Examples of individual psychometric functions and the corresponding PSE estimates are shown in Fig. S1. Prior to inferential statistical analyses, the distribution of the dependent measures in each condition was assessed for normality using the Shapiro–Wilk test. Depending on the normality of the data, paired t tests or Wilcoxon signed-rank tests were applied.

The speed of the AMR was computed from trajectory data obtained from the motion tracker mounted on the robot. The tracker provided three-dimensional position coordinates, from which the instantaneous speed was derived. The speed data for each trial were preprocessed in the following steps. First, outliers were excluded by removing data points exceeding ±3 standard deviations from the mean speed of the trial. Second, a median filter was applied to the remaining data to reduce noise.

For each trial, the mean speed was calculated during the constant-velocity phase of the approach. This phase was defined as the interval following the point of maximum acceleration, beginning when the instantaneous acceleration first fell below a near-zero threshold of 0.05 m/s^2^ and ending when it subsequently deviated from this threshold. Trialwise mean speeds were then averaged within each condition (7 speeds × 2 approach directions). Finally, these condition means were averaged across participants to obtain the grand mean speed for each condition.

The gaze behavior of the participants was quantified using data recorded by the head-mounted eye tracker, which provided gaze coordinates for each video frame during a trial. The center of the AMR was defined as the geometric center of the four corner points on the robot’s front surface, which were tracked using DeepLabCut [20]. For each frame, the Euclidean distance between the detected gaze point and the estimated center of the AMR was computed and used as a measure of gaze alignment.

## III. Results of Experiments 1 & 2

### A. Experiment 1

The PSEs for the two approach directions (front and behind) are shown in Fig. 2a. The mean PSEs were 0.73 ± 0.052 and 0.71 ± 0.036 for the front and behind conditions, respectively. A paired t test revealed a significant difference between approach directions (*t*(29) = 3.09, *p* < .01, Cohen’s *d* = 0.56). These results indicate that the AMR was perceived as moving faster when it was approaching from behind than when it was approaching from the front.

**Fig. 2.**
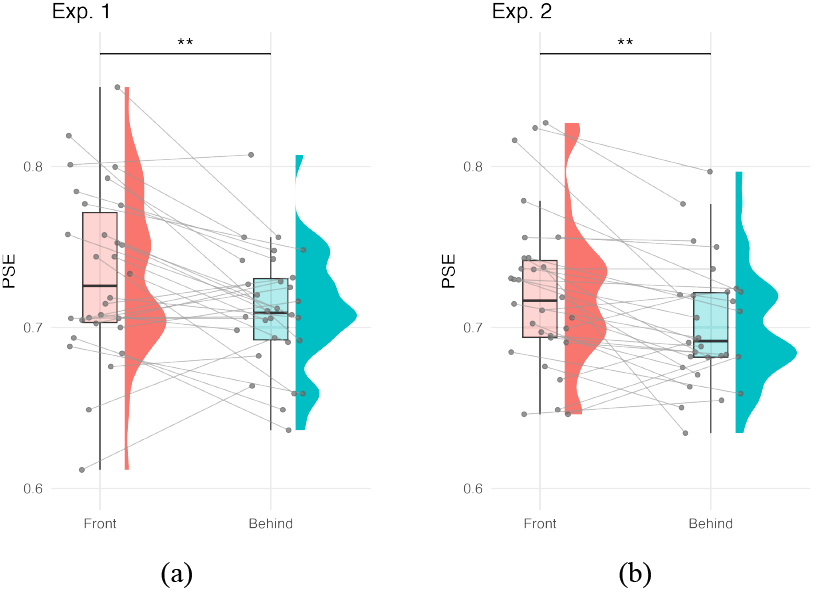
PSE results in Experiments 1 (a) and 2 (b). The horizontal axis represents the front and behind conditions. The vertical axis indicates the PSE, where a lower value signifies a greater perceived speed bias. The raincloud plot shows the individual data, box plot, and their distributions. Asterisks indicate significant differences: \*\**p* < .01.

In contrast, no significant difference was observed in the just noticeable difference (JND) between the front and behind conditions (*V* = 160, *p* = .22, *r* = 0.25). These findings indicate that the observed difference in perceived approach speed cannot be attributed to a difference in perceptual precision arising from viewing orientation (i.e., facing forward versus turning to look behind) but instead reflects a difference in perceptual bias. Notably, no significant condition-dependent differences in JND were observed in Experiments 2–5; therefore, the JND results are not reported to avoid redundancy.

To rule out the possibility that the effect was driven by unintended differences in the physical approach speed of the AMR, we computed the actual approach speeds for the front and behind conditions. Visual inspection of the speed profiles revealed no salient differences between approach directions across speed levels, suggesting that any residual variability in physical speed was negligible relative to the observed perceptual bias (Fig. S2a). Furthermore, to examine whether the effect could be explained by differences in gaze position between central and peripheral viewing, for each participant, we calculated the difference in (i) the mean distance between the gaze position and the center of the AMR and (ii) the PSE between the behind and front conditions. No significant correlation was observed between these two measures (*r*(28) = 0.17, *p* = .36; Fig. S3a). These results suggest that the observed difference in the PSE between conditions is unlikely to be explained by uncontrolled experimental artifacts. Rather, they indicate a genuine direction-dependent bias in perceived approach speed. Notably, this pattern of results was consistent across Experiments 1–3 (see Figs. S2 and S3).

### B. Experiment 2

Experiment 1 demonstrated that the AMR was perceived as moving faster in the behind condition than in the front condition. To examine whether this directional bias depended on the viewing direction used during the memorization of the reference speeds, Experiment 2 reversed the memorization direction such that participants learned the reference speeds while facing backward, whereas all the other procedures remained identical to those in Experiment 1.

The results of Experiment 2 are shown in Fig. 2b. A similar directional bias was observed, with the AMR again being perceived as moving faster in the behind condition than in the front condition. A paired t test confirmed a significant difference between conditions (*t*(29) = 3.44, *p* < .01, Cohen’s *d* = 0.63). These results indicate that the observed bias in perceived approach speed is independent of the viewing direction used during the memorization phase.

## IV. Methods of Experiment 3

The results of Experiments 1 and 2 consistently revealed that the perceived approach speed was higher in the behind condition than in the front condition, indicating that this directional bias is independent of the viewing direction used during the memorization phase. On the basis of these findings, Experiment 3 was designed to test the hypothesis that the observed bias arises from physiological constraints of the human visual system, specifically the binocular geometry.

When observers turn to look behind, the approaching AMR is viewed at an oblique angle relative to the head orientation, with the degree of obliqueness being proportional to the angle of body rotation (Fig. 3). As a result, the AMR in the behind condition is effectively perceived as approaching from a diagonal direction, in contrast to the direct frontal approach in the front condition. Diagonal motion induces interocular velocity differences (IOVDs), defined as discrepancies between the retinal image velocities in the two eyes [21], [22]. IOVDs constitute a primary cue for motion-in-depth perception, and neurophysiological studies have shown that neurons in the middle temporal (MT) area, which encode motion direction and velocity, are highly sensitive to IOVDs [23].

**Fig. 3.**
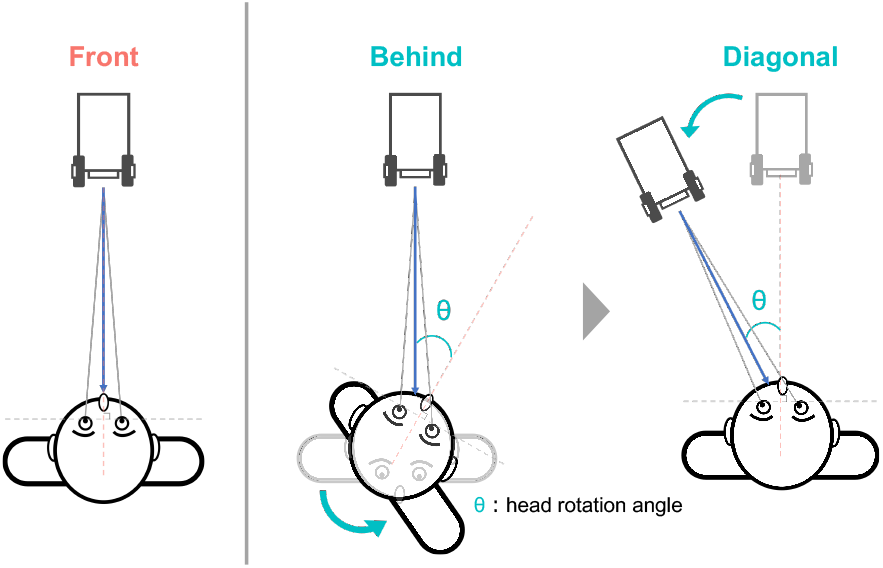
Illustration of the apparent position between front, behind, and diagonal conditions. The latter two conditions have the same visual inputs.

More specifically, as illustrated in Fig. 4a, the pattern of retinal image motion differs between the two conditions. In the front condition, retinal image motion is symmetric across the two eyes. In contrast, in the behind (diagonal) condition, the oblique approach induces interocular asymmetry such that the retinal image velocity is higher in one eye than in the other. This difference in interocular velocity may lead to an overestimation of approach speed, as the visual system weights the higher-velocity retinal signal more strongly during the perception of diagonal motion. Accordingly, Experiment 3 was designed to test whether the observed directional bias in perceived approach speed can be attributed to the IOVD.

**Fig. 4.**
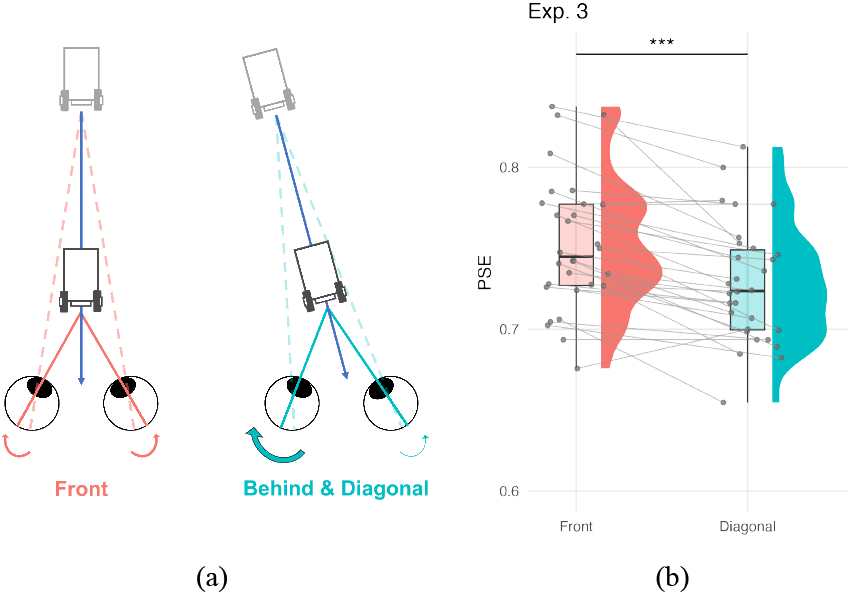
Schematic explanation of the IOVD for the both conditions and the results of Experiment 3. The right panel adopts the same format as that in Fig 2a does.

### A. Participants

Experiment 3 included 30 participants (9 females, 21 males, mean age 22.1 ± 1.4 years), matching the sample size used in Experiment 1.

### B. Apparatus

The apparatus was identical to that used in Experiment 1, except that the diagonal condition was employed instead of the behind condition.

### C. Procedure

The procedure was largely identical to that of Experiment 1, with the critical addition being a calibration phase conducted prior to the memorization task. During this phase, each participant’s head rotation angle (θ) when turning to look behind was measured. This angle was used to estimate the apparent position of the AMR relative to the observer’s viewpoint in the behind condition.

In the main task, perceived approach speed was compared between a frontal approach (front condition) and a diagonal approach originating from the estimated apparent position (diagonal condition; Fig. 3). The participants’ heads were stabilized using a chin rest to maintain a consistent forward-facing orientation. They were instructed to track the approaching AMR using eye movements only while keeping their head stationary throughout each trial. All the other aspects of the procedure were identical to those in Experiment 1.

### D. Data analysis

The data were analyzed in the same manner as were those in Experiment 1, except that the diagonal condition was treated as the behind condition. With respect to the gaze analysis, data from one participant were excluded because of a communication failure in the measurement equipment. Consequently, the gaze data from the remaining participants were used for the subsequent analysis.

## V. Results of Experiment 3

The PSEs for the front and diagonal conditions are shown in Fig. 4b. The mean PSEs were 0.75 ± 0.041 for the front condition and 0.73 ± 0.037 for the diagonal condition. A significant difference in PSE was observed between conditions (*t*(29) = 6.04, *p* < .001, Cohen’s *d* = 1.10), indicating that the diagonal approach was perceived as being faster than the frontal approach was. These results support the hypothesis that differences in interocular velocity contribute to direction-dependent bias in perceived approach speed.

## VI. Methods of experiments 4 & 5

The results of Experiments 1–3 suggested that the direction-dependent bias in perceived approach speed arises from IOVDs. To examine the robustness and generality of this effect, Experiments 4 and 5 were conducted in a virtual reality (VR) environment that replicated the physical setup of Experiment 1. In Experiment 4, participants wore a head-mounted display (HMD) that provided binocular visual input, allowing IOVDs to be reproduced under controlled visual conditions. We hypothesized that an approaching agent would be perceived as moving faster when approaching from behind than from the front, even in VR. Importantly, the virtual environment eliminated nonvisual cues associated with a physical robot, such as motor noise and airflow, enabling the assessment of whether the directional bias is driven primarily by visual information.

In Experiment 5, the visual appearance of the AMR was further simplified by replacing the robot model with a cylindrical object of the same size. This manipulation allowed us to test whether robot-specific appearance contributes to the observed bias in perceived approach speed.

### A. Participants

Experiments 4 and 5 were conducted as part of a university course. In Experiment 4, 40 students participated in the course, and data from 36 participants (7 females, 29 males; mean age = 21.2 ± 1.5 years) were included in the final analysis after absences and incomplete data were excluded. In Experiment 5, data from 30 participants (5 females, 25 males; mean age = 20.8 ± 0.6 years) were analyzed.

### B. Apparatus

The experimental stimuli were developed using Unity (2022.3.22f1) and SteamVR (2.14.5). Participants wore an HMD (HTC VIVE Cosmos) with a resolution of 1440 × 1700 pixels per eye and a refresh rate of 90 Hz. The virtual environment was designed to replicate the physical setting of Experiment 1.

In Experiment 4, the approaching agent was a three-dimensional model of the same AMR used in Experiment 1 (Fig. 5, left). The participant’s initial position was set at the origin of the virtual environment, and the AMR was programmed to approach from a distance of 4 m and stop 2 m from the participant. The velocity profile of the AMR was controlled to replicate the physical behavior observed in the real-world setting, consisting of a smooth acceleration phase, a constant-velocity phase at the target speed, and a gradual deceleration as the AMR approached the stopping point.

**Fig. 5.**
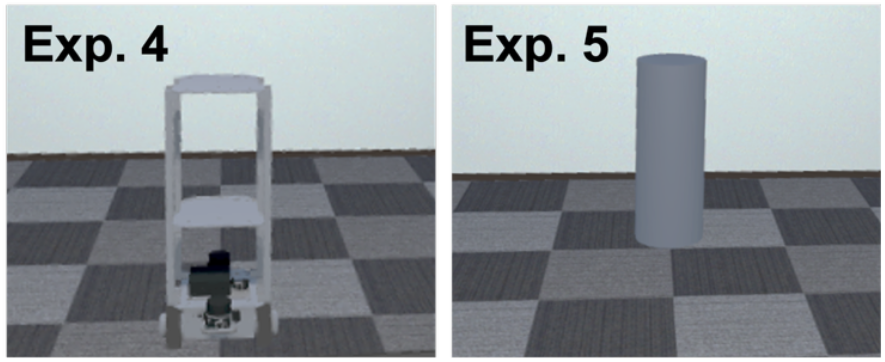
Stimuli used in Experiments 4 (AMR) and 5 (cylinder).

Experiment 5 was identical to Experiment 4, except that the AMR model was replaced with a primitive cylindrical object of the same size (Fig. 5, right). The cylinder dimensions and surface color were matched to those of the AMR to control for low-level visual properties.

### C. Procedure

The procedure was identical to that of Experiment 1.

### D. Data analysis

The data were analyzed in the same manner as in Experiment 1.

## VII. Result of Experiments 4 & 5

The PSEs for the two approach directions in Experiment 4 are shown in Fig. 6a. The mean PSEs were 0.74 ± 0.050 for the front condition and 0.72 ± 0.054 for the behind condition. A significant difference was observed between conditions (*t*(35) = 3.06, *p* < .01, Cohen’s *d* = 0.51), indicating that approach speed was perceived as being higher when the AMR approached from behind than when it approached from the front. These results demonstrate that direction-dependent perceptual bias is not restricted to physical environments and remains robust in a virtual setting. Moreover, because the virtual environment eliminated nonvisual cues such as motor noise and airflow, these findings suggest that the bias is driven primarily by visual information, which is consistent with an IOVD-based account. Similarly, Experiment 5 revealed a significant difference in PSEs between approach directions (*t*(29) = 5.16, *p* < .001, Cohen’s *d* = 0.94; Fig. 6b). These results indicate that the perceptual overestimation of approach speed for rearward approaches persists even when the approaching agent is a simple geometric object, suggesting that the effect is independent of robot-specific appearance or visual complexity.

**Fig. 6.**
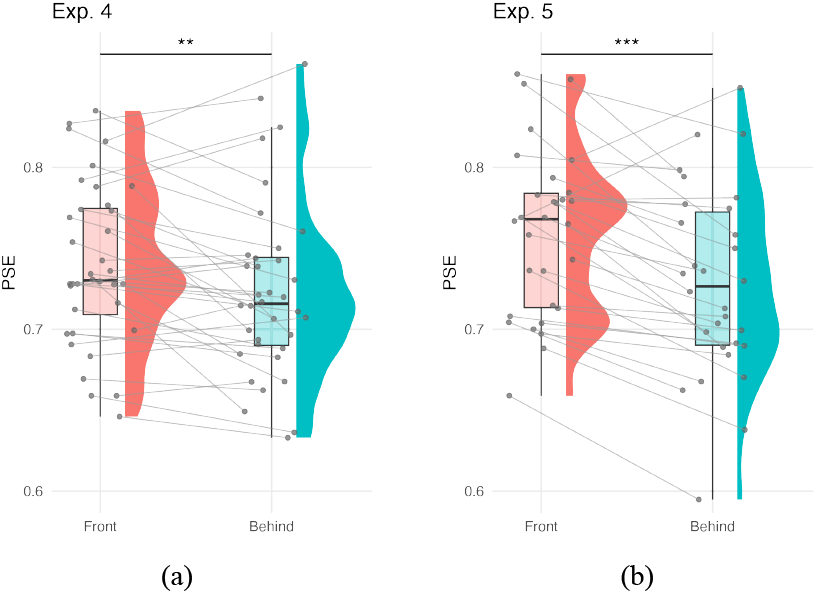
PSE results obtained in Experiments 4 (a) and 5 (b). The format was the same as that of Experiment 1.

## VIII. Discussion

Our results consistently demonstrate that approach speed is perceived as being higher when an object approaches from behind than when it approaches from the front across a range of experimental settings. The robustness of this effect indicates that the bias is rooted in a fundamental physiological mechanism, specifically IOVDs, as directly tested in Experiment 3, rather than in situational factors such as viewing orientation during reference-speed learning or spatial context (Experiments 1 and 2), nonvisual cues including motor noise (Experiment 4), or object appearance (Experiment 5).

How do interocular velocity differences operate within the visual system to yield the observed bias? In the neural processing of visual motion, signals initially detected in the primary visual cortex (V1) are integrated into the middle temporal (MT) area to estimate three-dimensional velocity [24]. Previous studies have shown that approximately half of MT neurons are highly sensitive to IOVD cues [23]. Within this computational framework, we propose that the visual system exhibits a perceptual bias toward the higher-velocity signal when retinal inputs are asymmetric across the two eyes. This bias provides a parsimonious account of our findings because the diagonal gaze required in the behind condition inherently generates IOVDs, leading to a systematic overestimation of approach speed across a wide range of experimental settings.

In addition to visual–mechanistic factors, psychological threat may also contribute to the overestimation of approach speed from behind. Neuhoff reported that a wide range of organisms, from insects to humans, exhibit defensive responses to visually looming objects and that such stimuli in humans activate the autonomic and motor systems in preparation for avoidance behavior [25]. An approach from behind may be implicitly interpreted as a chasing scenario, thereby amplifying perceived speed through a biologically grounded defensive response. This interpretation is consistent with findings by Balcetis, who showed that perceptual biases, particularly in distance perception, can serve adaptive functions by facilitating rapid self-defensive actions in potentially threatening situations [26]. These observations suggest that the human perceptual system may be biased to overestimate approach speed during head turning as an adaptive strategy that supports rapid responses to threats originating from the rear.

Furthermore, previous studies have reported that approaches from behind elicit greater psychological discomfort than approaches from the front do [14]. Our findings suggest that the overestimation of approach speed for rearward approaches may contribute to this heightened discomfort. Notably, the results of Experiment 5 revealed that this overestimation occurred regardless of object type, implying that the associated discomfort reflects a general perceptual phenomenon rooted in human sensory processing rather than a response to robot-specific visual appearance.

Crucially, these results highlight a fundamental asymmetry in perception between the front space and rear space, underscoring the need for asymmetric motion design in human–robot interactions. For example, autonomous delivery robots operating in pedestrian environments should incorporate control strategies that either avoid approaching humans from behind or substantially reduce their approach speed when rearward approaches are unavoidable. Such perception-aware control strategies can mitigate the psychological impact of robot motion by compensating for the human tendency to overestimate approach speed from behind.

This study has several limitations. First, although we quantified a robust perceptual bias in approach speed, we did not directly measure the associated emotional responses, such as perceived threat or subjective comfort. While our results indicate that speed perception itself is driven primarily by physiological factors, affective evaluations of robot motion are inherently multifaceted. Because approaches from behind are generally experienced as more unpleasant than frontal approaches are [14], the influence of robot appearance and motion may differ when assessed in terms of emotional comfort rather than purely sensory perception.

Second, optimal robot behavior is highly context dependent. While lower approach speeds may be desirable for delivery robots operating in public pedestrian environments to promote social acceptance, collaborative robots in industrial settings may require higher speeds to maintain operational efficiency. Moreover, real-world human–robot interactions encompass a wide range of approach angles, environmental constraints, and social contexts that were not fully captured in our controlled experimental design. To address these limitations, future work will focus on developing perception-aware control algorithms that explicitly incorporate the present findings. We aim to evaluate the effectiveness of such algorithms in complex, real-world pedestrian environments, with the goal of increasing both safety and social acceptance while balancing physical efficiency and human psychological comfort across diverse interaction scenarios.

## IX. Conclusion

This study examined direction-dependent differences in the perceived approach speed of an AMR. Across a series of five experiments, we consistently found that approaches from behind are perceived as being faster than frontal approaches are. Our results indicate that this perceptual bias is primarily attributable to interocular velocity differences arising from oblique visual geometry during head turning, reflecting a fundamental characteristic of the human visual system. These findings underscore the importance of incorporating asymmetries in human perception into robot motion design and provide empirical motivation for perception-aware and direction-sensitive control strategies in human–robot interaction.

## Notes

https://osf.io/fsb8t/

### Competing Interest Statement

The authors have declared no competing interest.

### Funding

- Japan Society for the Promotion of Science: JP25K21323, JP25H01141, JP23KK0183
- Foundation of Public Interest of Tatematsu
- Amano Institute of Technology

