## Supplementary Information for "Direction-dependent bias in the perceived approach speed and its implications for robot motion design"

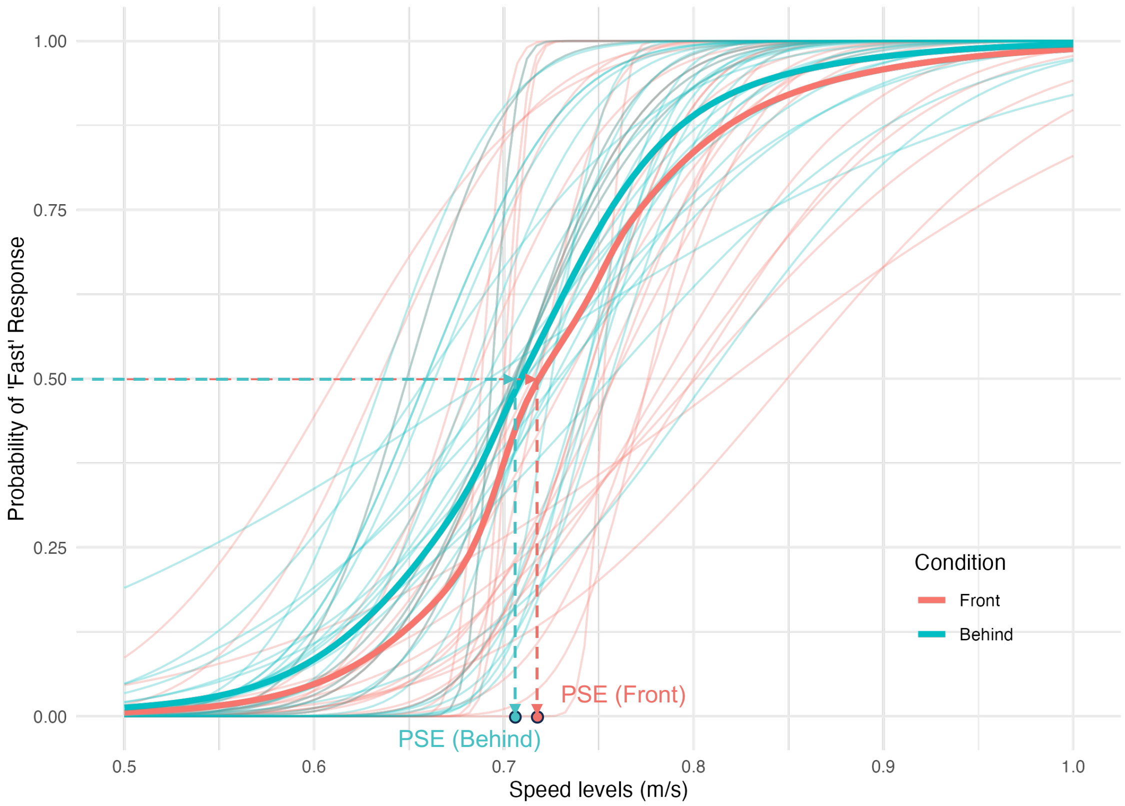


**Fig. S1.** Psychometric curves for the front and behind conditions in Experiment 1. The horizontal axis represents the AMR's speed. The vertical axis indicates probability of the response being "fast". Thin and thick lines represent individual participant data and group averages, respectively, for both approach conditions. The points at the end of the dashed lines indicate the PSE for each approach condition.

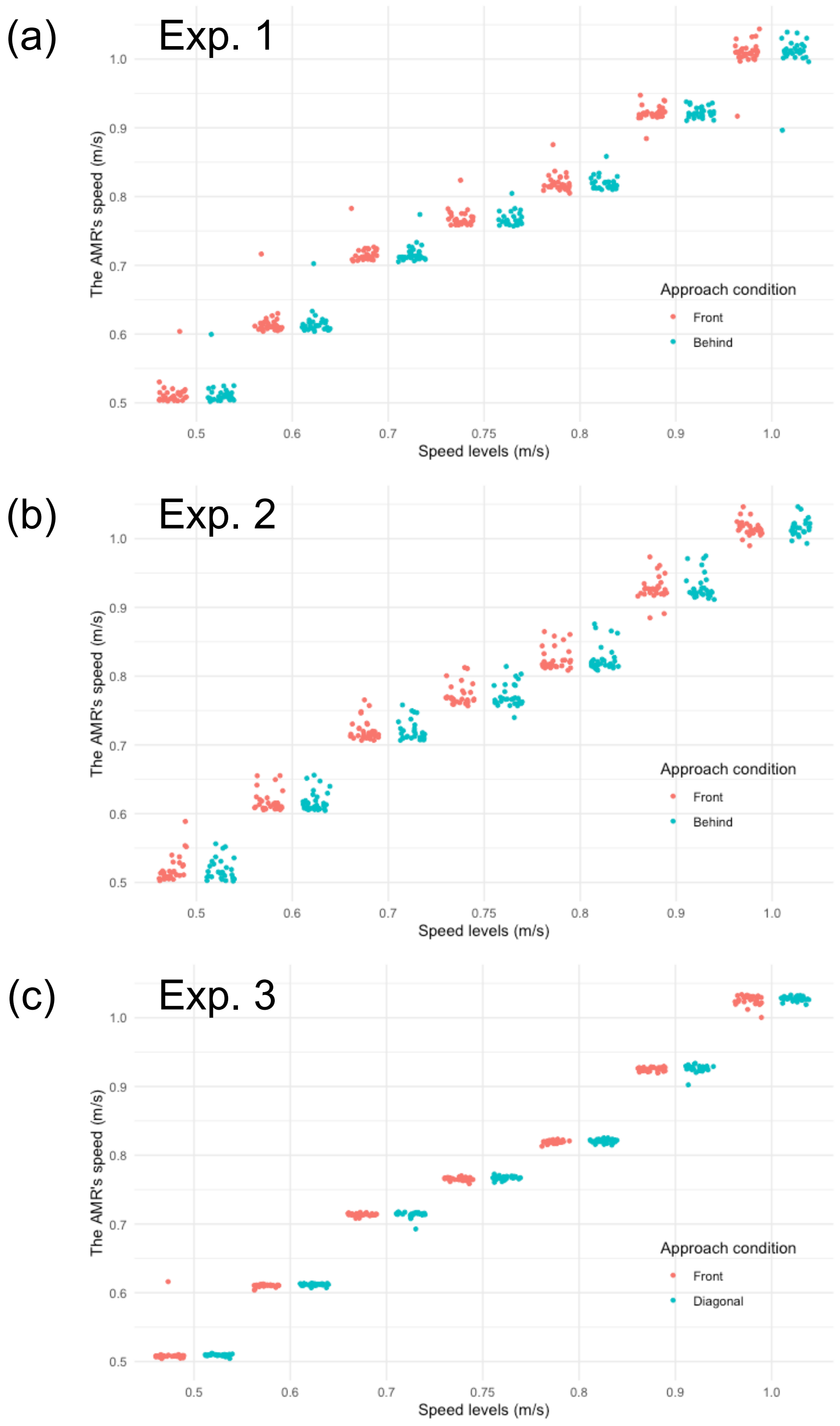


**Fig. S2.** Actual speed of the AMR in the front and behind conditions. The horizontal axis represents the seven speeds. The vertical axis indicates the actual speed of the AMR. Each point represents the individual data. Panels (a), (b), and (c) illustrate the results of Experiments 1, 2, and 3, respectively.

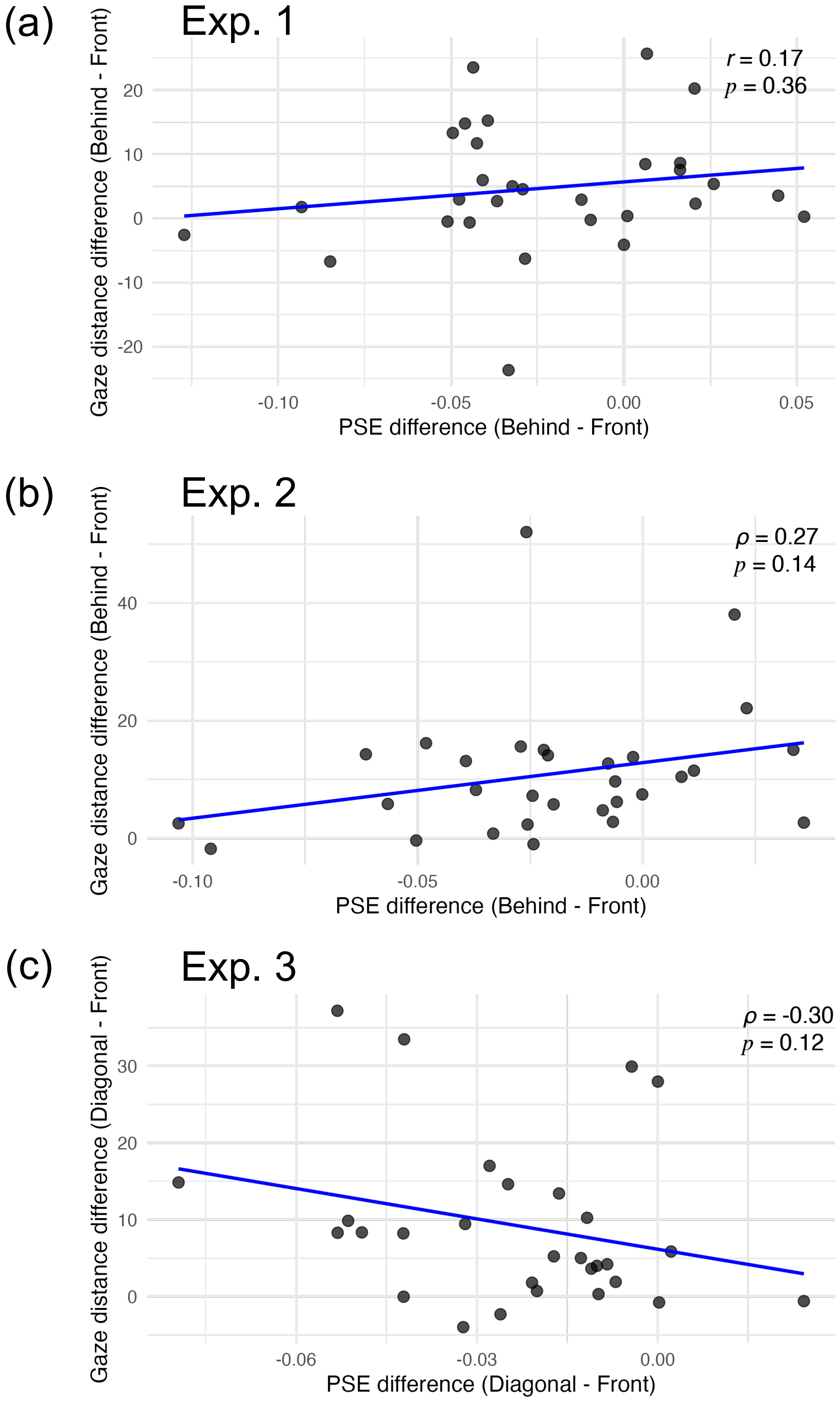


**Fig. S3.** Correlation between the difference in PSE and the difference in the gaze distance for both approach conditions. The horizontal axis represents the difference in PSE (behind/diagonal - front). The vertical axis indicates the difference in gaze distance (behind/diagonal - front). The points show the individual data. Panels (a), (b), and (c) illustrate the results of Experiments 1, 2, and 3, respectively.
